# Medial temporal atrophy in preclinical dementia: visual and automated assessment during six year follow-up

**DOI:** 10.1101/2020.03.05.979229

**Authors:** Gustav Mårtensson, Claes Håkansson, Joana B. Pereira, Sebastian Palmqvist, Oskar Hansson, Danielle van Westen, Eric Westman

## Abstract

Medial temporal lobe (MTL) atrophy is an important morphological marker of many dementias and is closely related to cognitive decline. In this study we aimed to characterize longitudinal progression of MTL atrophy in 93 individuals with subjective cognitive decline and mild cognitive impairment followed up over six years, and to assess if clinical rating scales are able to detect these changes. All MRI images were visually rated according to Scheltens’ scale of medial temporal atrophy (MTA) by two neuroradiologists and AVRA, a software for automated MTA ratings. The images were also segmented using FreeSurfer’s longitudinal pipeline in order to compare the MTA ratings to volumes of the hippocampi and inferior lateral ventricles. We found that MTL atrophy rates increased with CSF biomarker abnormality, used to define preclinical stages of Alzheimer’s Disease. Both AVRA’s and the radiologists’ MTA ratings showed a similar longitudinal trajectory as the subcortical volumes, suggesting that visual rating scales provide a valid alternative to automatic segmentations. While the MTA scores from each radiologist showed strong correlations to subcortical volumes, the inter-rater agreement was low. We conclude that the main limitation of quantifying MTL atrophy with visual ratings in clinics is the subjectiveness of the assessment.

## 1 Introduction

Atrophy of the medial temporal lobe (MTL) is an important diagnostic biomarker in many different dementias, including Alzheimer’s Disease (AD). In research we quantify atrophy using automatic softwares that compute volume or thickness measures of regions of interests, specified by a neuroanatomical atlas. These softwares are either not sufficiently reliable for clinical usage, and the ones that have been approved are not widely implemented. To quantify atrophy in clinics, radiologists visually assess the degree of atrophy in a brain region according to established rating scales.

The most widely used rating scale in clinical practice is Scheltens’ scale of Medial Temporal Atrophy (MTA) (Scheltens et al., 1992; Vernooij et al., 2019). The MTA scale quantifies the level of atrophy in hippocampus (HC) and its surrounding structures, the choroid fissure and inferior lateral ventricle (ILV). The MTA scale has been shown to reliably distinguish individuals with AD from healthy elderly (Scheltens et al., 1992; Wahlund et al., 1999; Westman et al., 2011). It is an ordinal scale ranging from 0 (no atrophy) to 4 (end-stage atrophy) where an integer score is given for each hemisphere. In Fig. 1 we provide examples of each score. Several studies have reported on the diagnostic ability and relevant clinical cut-offs of the MTA scale (Westman et al., 2011; Scheltens et al., 1992; Ferreira et al., 2015), and others have argued for the importance of reporting MTA in the clinical routing (Torisson et al., 2015; Håkansson et al., 2019; Wahlund et al., 2017).

**Figure 1:**
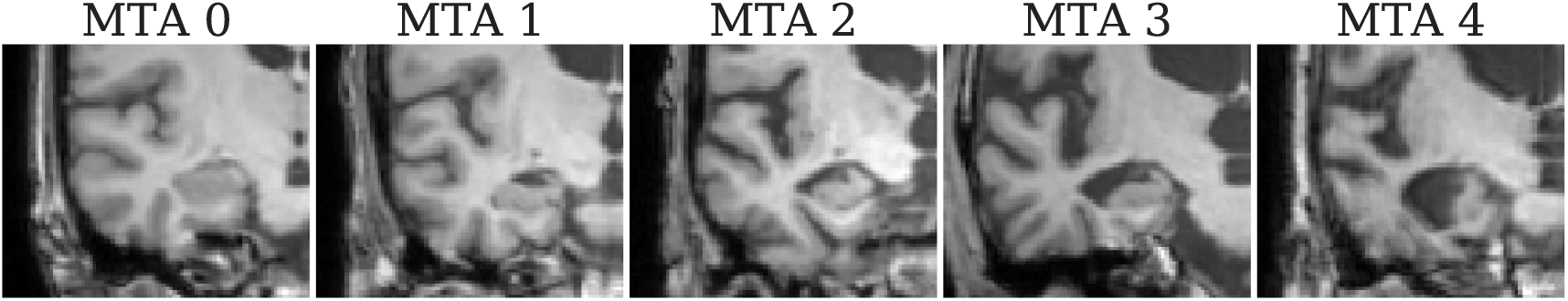
Example of the Scheltens’ MTA scale, with progressive atrophy of the hippocampus, the choroid fissure and the inferior lateral ventricle. The image selected for each score was given the same rating by both radiologists in this study. Each hemisphere is rated individually.

Longitudinal progression of medial temporal atrophy, quantified through e.g. hippocampal volumes, has been studied in cognitively normal subjects as well as in preclinical, prodromal and probable AD (Rusinek et al., 2003; Ridha et al., 2006; Henneman et al., 2009a; Pettigrew et al., 2017). The reported annual decrease in HC volume varies between the studies. Rusinek et al. (2003) found a 0.36% volume loss/year for cognitively stable subjects, and a greater loss (1%/year) in individuals with cognitive decline. Henneman et al. (2009a) reported 2.2% annual HC volume loss for healthy controls, with greater atrophy rates in patients with MCI (−3.8%/year) and AD (−4.0%/year). Another study reported an up to 8% decrease in HC volume per year in asymptomatic individuals at risk of familial AD (Fox et al., 1996). Despite the large interest in longitudinal MTL atrophy, no studies have investigated how these corresponds to clinical MTA ratings.

The aim of this paper is to investigate longitudinal changes in the MTL in individuals with subjective cognitive decline (SCD) or mild cognitive impairment (MCI), and whether clinical rating scales can detect these changes. Two neuroradiologists and AVRA (Automatic Visual Ratings of Atrophy)—our recently developed software providing automated continuous MTA scores—rated 93 individuals scanned four times over six years using Scheltens’ MTA scale. Further, all images were segmented using the longitudinal FreeSurfer pipeline to extract hippocampal and inferior lateral ventricle volumes. We calculated atrophy rates of the MTL for visual and automated measures to understand what progression to expect in different stages of preclinical dementia.

## 2 Methods

### 2.1 Study population

The study population was part of the prospective and longitudinal Swedish BioFINDER (Biomarkers For Identifying Neurodegenerative Disorders Early and Reliably) study (www.biofinder.se) and comprised non-demented people with subjective and objective cognitive decline. All patients were consecutively enrolled at three outpatient memory clinics and were assessed by physicians specialized in dementia disorders. Inclusion criteria were: 1) referred to the memory clinics because of cognitive symptoms, 2) not fulfilling dementia criteria, 3) MMSE score of 24–30 points, 4) age 60–80 years and, 5) fluent in Swedish. Exclusion criteria were: 1) cognitive impairment that without doubt could be explained by a condition other than prodromal dementia, 2) severe somatic disease, and 3) refusing lumbar puncture or neuropsychological investigation. A neuropsychological battery assessing four broad cognitive domains including verbal ability, visuospatial construction, episodic memory, and executive functions was performed and a senior neuropsychologist then stratified all patients into those with SCD (no measurable cognitive deficits) or MCI according to the consensus criteria for MCI (Petersen, 2004). From this larger cohort we selected all individuals who had been followed up three times over the course of six years.

As in the work by Pettigrew et al. (2017), we stratified the cohort based on abnormality in CSF amyloid-*β* (A) and phosphorylated tau (T) levels analyzed with Euroimmun essays (EUROIMMUN AG, Lübeck, Germany). We applied the cut-off A*β*_42_/A*β*_40_ < 0.10 (Janelidze et al., 2016) to define A^+^ and p-tau > 72 pg/ml (Mattsson et al., 2018) for T^+^. This yielded the subgroups A^−^T^−^ (i.e. denoting normal amyloid-*β* and p-tau levels), A^+^T^−^, and A^+^T^+^. No individuals displayed the CSF combination A^−^T^+^. Demographics and clinical characteristics of these groups are summarized in Table 1.

**Table 1:** Demographics of the included participants at baseline. *p*-values were computed using Kruskal-Wallis H-test, testing the null hypothesis that medians are equal in all subgroups.

| | All | A<sup>-</sup>T<sup>-</sup> | A<sup>+</sup>T<sup>-</sup> | A<sup>+</sup>T<sup>+</sup> | $p$-value |
| --- | --- | --- | --- | --- | --- |
| $N$ | 93 | 54 | 18 | 21 | — |
| SCD/MCI | 61/32 | 42/12 | 8/10 | 11/10 | — |
| Age at bl. | 70.06 ± 5.41 | 69.71 ± 5.57 | 70.18 ± 4.55 | 70.86 ± 5.58 | 0.208 |
| Sex, F (%) | 57.0 | 64.8 | 50.0 | 42.9 | 0.001 |
| ApoE4 carriers (%) | 38.7 | 14.8 | 66.7 | 76.2 | <.001 |
| Education (years) | 12.01 ± 3.30 | 11.91 ± 3.34 | 11.67 ± 3.42 | 12.57 ± 3.02 | 0.108 |
| MMSE at bl. | 28.26 ± 1.72 | 28.57 ± 1.46 | 28.06 ± 1.75 | 27.62 ± 2.06 | <.001 |
| ADAS-DWR at bl | 4.24 ± 2.73 | 3.41 ± 2.39 | 5.17 ± 2.41 | 5.38 ± 3.05 | <.001 |
| CSF A$\beta_{42/40}$ ratio | 0.12 ± 0.04 | 0.15 ± 0.03 | 0.09 ± 0.02 | 0.07 ± 0.02 | — |
| CSF A$\beta_{42}$ (pg/ml) | 611.8 ± 259.4 | 788.0 ± 182.7 | 370.6 ± 135.5 | 399.3 ± 145.3 | — |
| CSF p-tau (pg/ml) | 56.7 ± 36.5 | 35.7 ± 10.9 | 47.9 ± 14.4 | 113.4 ± 27.6 | — |
| $N$ conv. to dementia (to AD) | 19 (13) | 5 (0) | 5 (5) | 9 (8) | — |

### 2.2 MRI protocol

All T_1_-weighted MRI scans were acquired with an MPRAGE protocol on a 3T Siemens TrioTim with the following parameters: 1.2 mm slice thickness, 0.98 mm inplane resolution, 3.37 ms Echo Time, 1950 ms Repetition Time, 900 ms Inversion Time, and 9° Flip angle.

### 2.3 Visual assessments

Two neuroradiologists (*Rad. 1* and *Rad. 2*) rated all available images according to Scheltens’ MTA scale (Scheltens et al., 1992), see Fig. 1. The raters were blinded to sex, age, diagnosis, amyloid-*β* and tau status, subject ID and timepoint to not bias the ratings. Both radiologists assess MTA on a regular basis as part of their clinical work but have not trained together to facilitate ratings consistency.

### 2.4 Automated methods

For automated MTA ratings we used our recently proposed software AVRA^1^ v0.8 (Mårtensson et al., 2019a). Briefly, AVRA is a deep learning model that was trained on more than 3000 MRI images from multiple cohorts rated by a single radiologist (none of the raters in the current study). It is based on convolutional neural networks and predicts MTA from features extracted from the raw images (i.e., not volumetric data), similar to how a radiologist would perform the assessment. The model has previously demonstrated substantial inter-rater agreement levels in multiple imaging cohorts from various memory clinics (Mårtensson et al., 2019a,b). The first step of the processing pipeline of AVRA is to align the anterior and posterior commissures (AC-PC) using FSL FLIRT (Jenkinson and Smith, 2001; Jenkinson et al., 2002). We visually inspected the rigid registrations to ensure that the AC-PC alignment had not failed, but no AVRA ratings were discarded based on this. Contrary to the radiologist ratings, AVRA outputs continuous MTA scores. This allows for capturing more subtle longitudinal changes in the MTA scores that is not possible with a discretized scale.

All MRI scans were processed through TheHiveDB system (Muehlboeck et al., 2014) with FreeSurfer^2^ 6.0.0 for automatic segmentation of cortical and subcortical structures, such as hippocampi and inferior lateral ventricles (Dale et al., 1999; Fischl et al., 2004). First, all images were processed cross-sectionally, and each output was visually inspected to detect images with inaccurate hippocampal or ventrical segmentation. Images that passed quality control were re-processed with FreeSurfer’s longitudinal pipeline for more consistent segmentation (Reuter et al., 2012). The longitudinal output was once again visually inspected and cases with poor hippocampal or ventrical segmentations excluded. In total, 339 (out of 372) images from 87 (out of 93) participants were included in the study for further analyses.

### 2.5 Analyses

The analyses revolve around two central themes: to study the sensitivity and reliability of MTA ratings in a longitudinal setting, and to characterize medial temporal atrophy in preclinical dementia.

For the first theme we used Cohen’s weighted kappa *κ_w_* ∈ [−1,1] to assess the agreement between two sets of ratings. As there is no ground truth available, *κ_w_* is a common metric to report in studies using visual ratings, where a high inter- and intra-rater agreement suggests that the rater is reliable and consistent. We further compared the manual and automated ratings to hippocampal and inferior lateral ventricle volumes. Although MTA is rated in a single slice—and does not assess volumes—we assume that reliable ratings should (anti-)correlate strongly with HC and ILV volumes. We studied visual rating sensitivity, i,e. their ability to capture MTL atrophy, by comparing within-subject changes in MTA ratings (“ΔMTA”) to changes in HC and ILV volume (“ΔHC” and “ΔiLV”).

Characterizing MTL atrophy in preclinical dementia was done by studying the cross-sectional and longitudinal progression of MTA scores, HC volumes and ILV volumes as a function of age. We approximated the average annual change in MTA scores (“ΔMTA/year”), Mini Mental State Examination (MMSE), Alzheimer’s Disease Assessment Scale-delayed word recall (ADAS-DWR), HC and ILV volumes by fitting a least-squares regression line for each individual and measure. To study the effect of clinical status (i.e. SCD or MCI), we performed additional analyses on SCD and MCI subjects separately within each CSF group. The analyses including volumetric data were performed on the subset of images that passed the visual quality control.

## 3 Results

The rating agreements between radiologists and AVRA, and their correlations to HC and ILV volumes, are shown in Table 2. The weighted kappa agreements between the raters ranged from *fair* (*κ_w_ ∈* [0.2,0.4)) to *substantial (κ_w_ ∈* [0.6,0.8)) (Landis and Koch, 1977). All sets of ratings demonstrated similar Spearman correlations strengths to HC (*r_s_ ∈* [-0.57,-0.47]) and ILV (*r_s_ ∈* [0.78,0.88]) volumes. Violinplots illustrating the distribution of HC and ILV volumes per MTA score and rater are shown in Fig. 2, where we note that Rad. 1 systematically gave higher MTA scores than Rad. 2. We include confusion matrices between rating sets for left and right hemispheres as Supplementary data, Tables S1–S3.

**Table 2:**
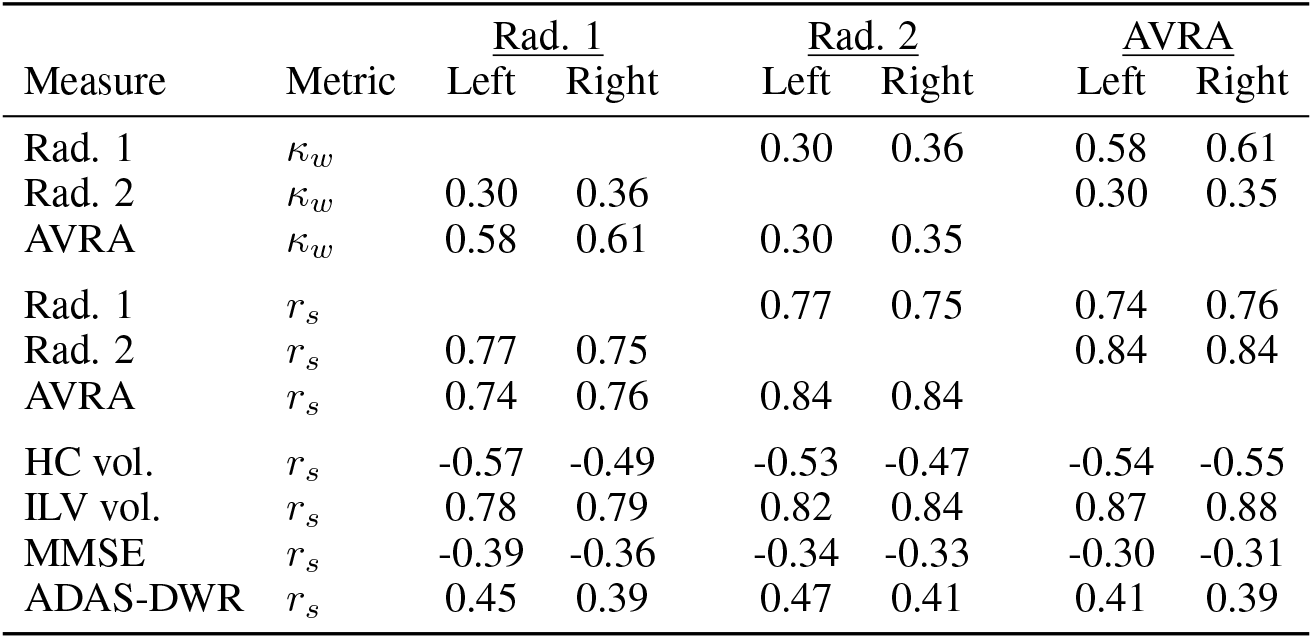
Inter-rater agreements (*κ_w_*) and Spearman correlations (*r_s_*) between radiologists’ ratings, hippocampal (HC) and inferior lateral ventricle (ILV) volumes.

**Figure 2:**
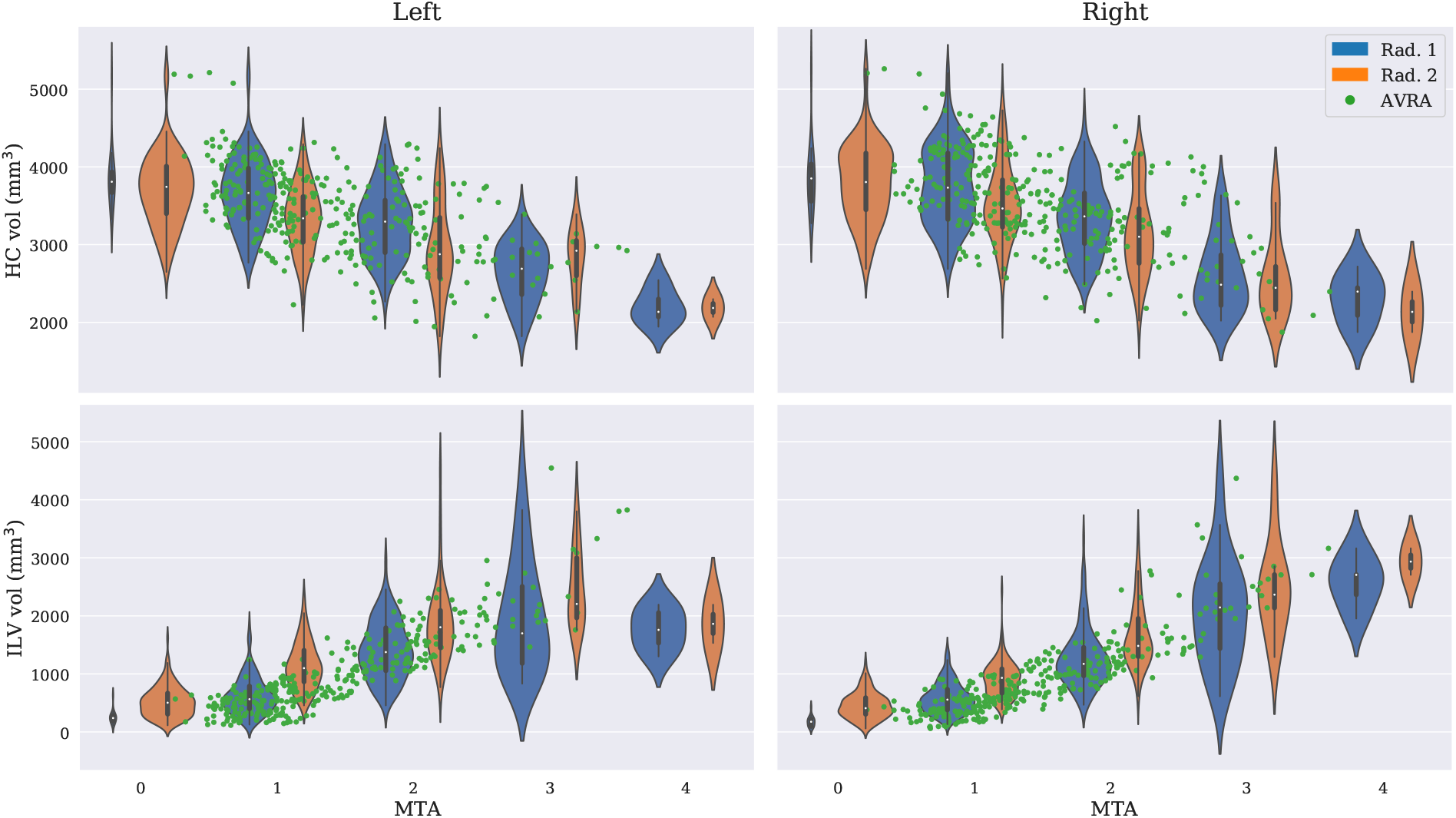
Violinplots of the radiologists’ MTA ratings and corresponding hippocampal volume (*top*) and inferior lateral ventricle volume (*bottom*). The width of the violins shows the distribution over volumes for each rating and rater, and the area indicates the number of images given a specific rating. The green dots show AVRA’s MTA rating for each image.

In Table 3 the baseline characteristics and average annual progression rates among study participants for all sets of ratings, MTL volumes, MMSE and ADAS-DWR are shown. No clear pattern was found between CSF groups in the cross-sectional baseline measures. However, all MTL measures showed that the atrophy rates increased with progressing AD CSF pathology, with the exception of the ratings from Rad. 1 that showed a milder progression in the A^+^T^−^ group. By assessing the SCD and MCI patients separately, we observed that the CSF group differences in atrophy rates were larger in the MCI subset. We further noted that the atrophy rates were greater in the SCD subjects in A^+^T^+^ than in the MCI patients in the A^−^T^−^ group.

**Table 3:**
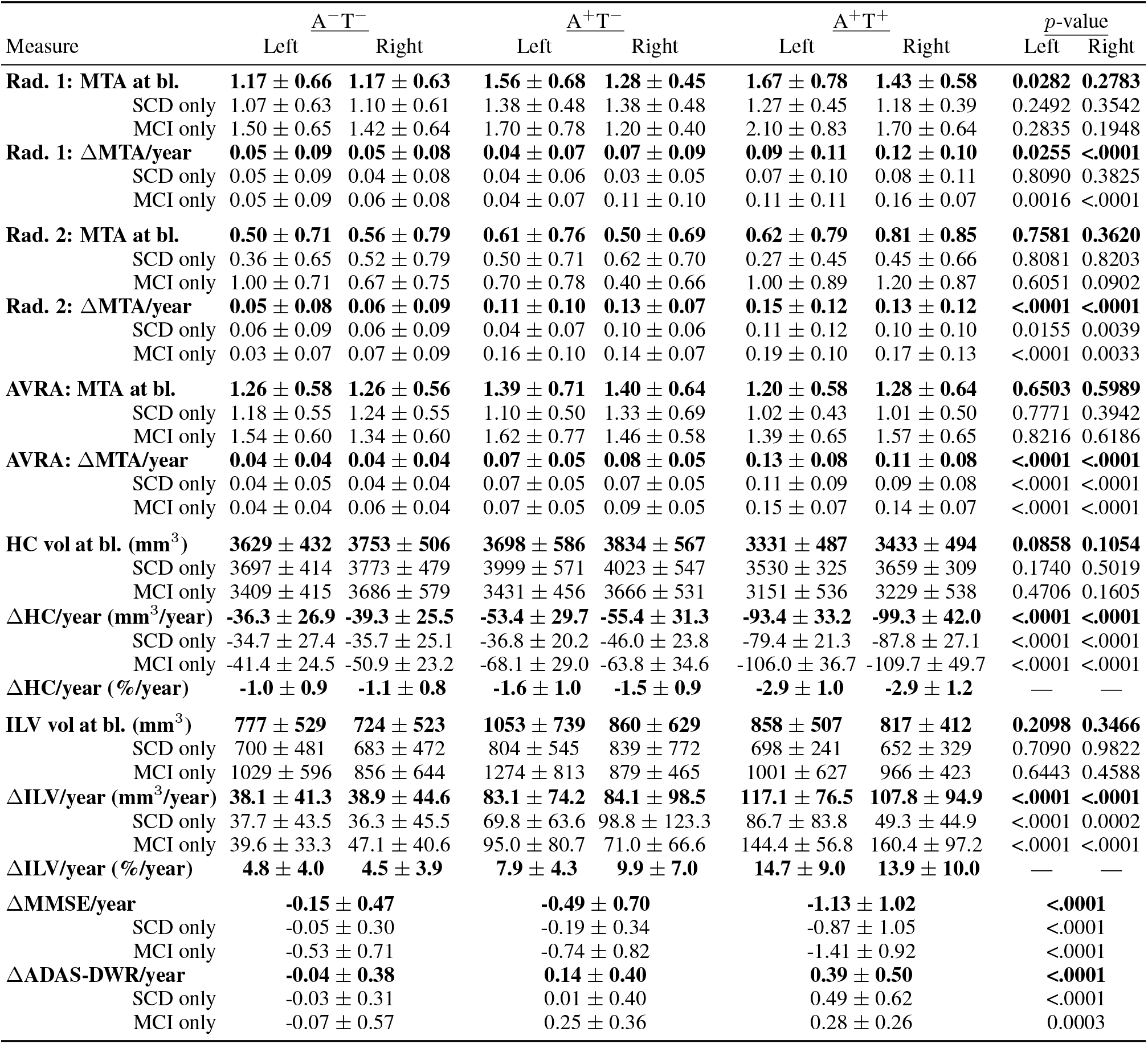
Average baseline (bl) MTA ratings and volumes, and average annual change for individuals with different CSF statuses. Rows in bold denotes entries where the whole CSF group was considered (i.e. SCD’s and MCI’s), and ‘SCD/MCI only’ refers to the subset of SCD/MCI subjects within the CSF group. ΔMTA/year refers to the average annual change in MTA score of the study participants. The reported p-values were computed using Kruskal-Wallis H-test to test the null-hypothesis that the population medians of all groups were equal. Applying a Bonferroni correction to a significance level of *α* = 0.01 would render all p-values <.0001 significant.

In Fig. 3 the trajectories of each study participant are displayed for left MTA (predicted with AVRA), HC volume and ILV volume respectively. (The measures of the right hemisphere showed similar characteristics, and are provided as Supplementary data). The longitudinal trajectories of the FreeSurfer measures were generally smoother than AVRA’s MTA scores, which were not monotonically increasing for individuals, suggesting some degree of rating variability. From Fig. 3 we see that the MTL measures of the MCI patients (orange lines) were generally more pathological than the SCD subjects (blue lines), which is confirmed in Table 3. We include examples of MRI scans for all timepoints for randomly selected participants as Supplementary data, Fig. S1.

**Figure 3:**
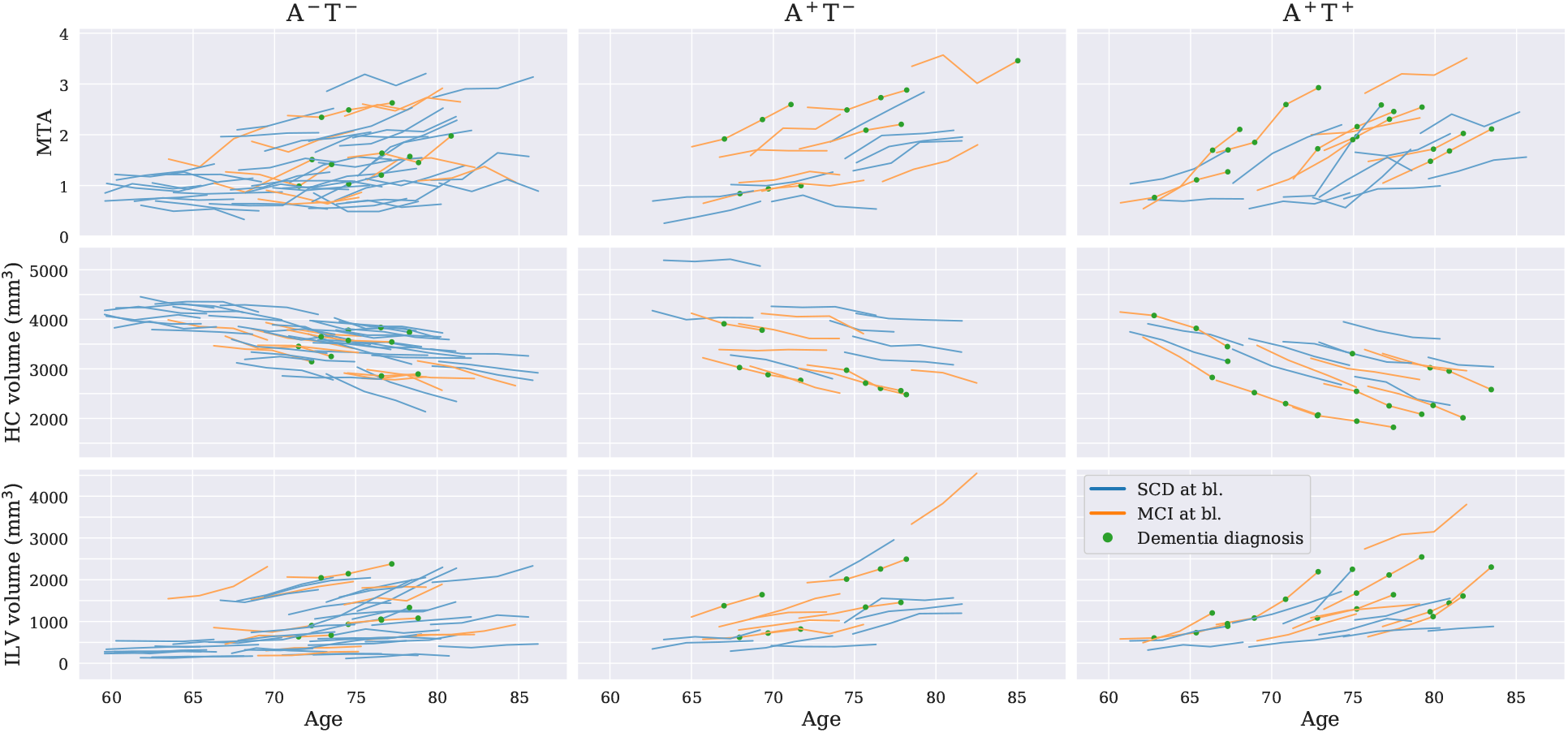
*Top*: AVRA’s left MTA ratings plotted against age at scan time for different combinations of *Aβ* and p-tau abnormality. *Middle/bottom:* corresponding plots for left hippocampal (HC)/inferior lateral ventricles (ILV) volumes. Orange and blue lines show individual trajectories for SCD and MCI patients, respectively. The green dots show if a patient was diagnosed with dementia at the given timepoint.

To study the sensitivity of the discrete radiologist ratings, we investigated the changes in MTA scores and MTL volumes compared to baseline. In Fig. 4 we show kernel density plots that estimate the distribution of ΔHC and ΔILV for follow-up images given the same MTA score (ΔMTA=0), +1 MTA (ΔMTA=1) and +2 MTA (ΔMTA=2). Both radiologists show similar distributions for ΔMTA=0 and the ΔMTA=1 entries, with a larger shift in means for ΔMTA=2. From these results it was possible to estimate that when ΔHC equaled −238 mm^3^ (−8%) and −235 mm^3^ (−7%) mm^3^ it became more likely that the image was being rated with a higher MTA score, for Rad. 1 and Rad. 2 respectively. Corresponding values for ΔILV were 225 mm^3^ (27%) and 254 mm^3^ (33%).

**Figure 4:**
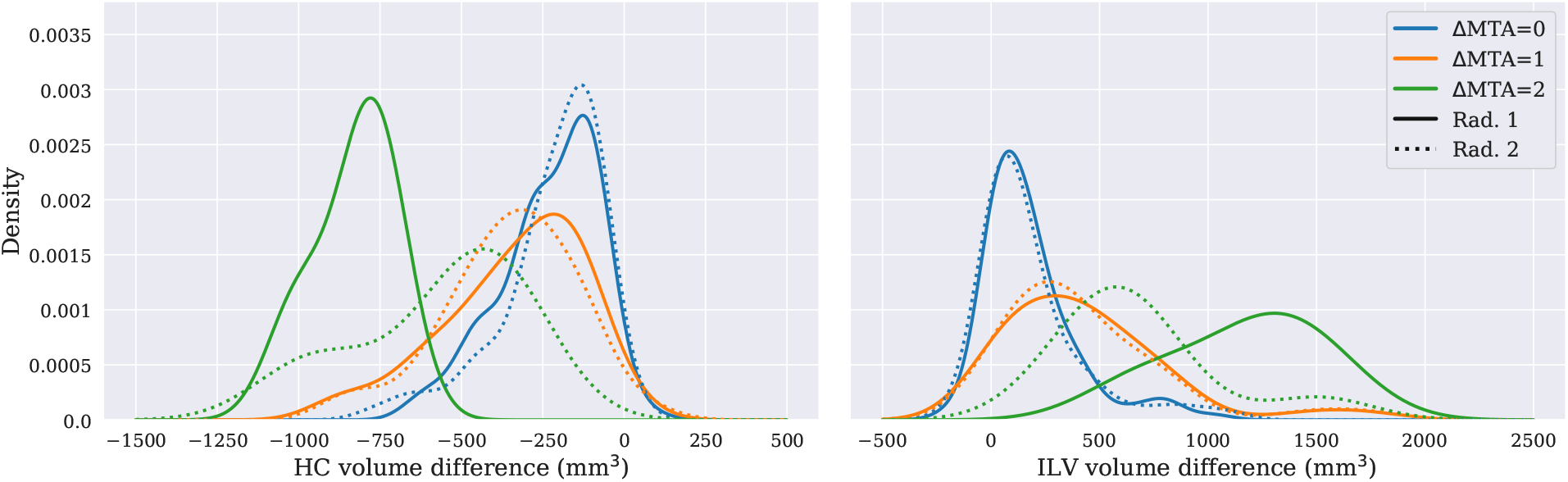
Shows distribution (kernel density plots) of the change in HC (left) and ILV (right) volumes between baseline and follow-up scan. A follow-up image rated the same as the baseline scan are in blue (“0 MTA”), 1 MTA score higher (“+1 MTA”) in orange, and 2 MTA scores higher (“+2 MTA”) in green. Solid lines are ratings from Rad. 1 and dotted lines from Rad. 2.

## 4 Discussion

In this study we investigated longitudinal medial temporal atrophy in preclinical dementia, and to what extent it is possible to capture these changes with the Scheltens’ MTA scale. We found that both radiologists provided reliable ratings, capable of capturing longitudinal progression, despite low inter-rater agreement. This was due to systematic rating differences between the radiologists, which highlights the issue of using subjective methods to quantify atrophy. Further, we observed increased MTL atrophy rates with worsening cognition and CSF AD pathology. This is the first study to investigate longitudinal MTL atrophy using MTA ratings, which helps bridge the gap between neuroimaging research and clinical radiology.

The rating agreement was only moderate between Rad. 2 and Rad. 1, as well as between Rad. 2 and AVRA. This is slightly lower than inter-rater agreements reported in studies using MTA, normally in the range *κ_w_ ∈* (0.6, 0.9) (Koedam et al., 2011; Cavallin et al., 2012b; Velickaite et al., 2017; Ferreira et al., 2017). All sets of ratings showed strong correlation to both HC and ILV volumes. This was reasonable, given that another recently proposed model estimating MTA was based on a linear combination of HC and ILV volumes (Koikkalainen et al., 2019). Our reported Spearman correlations between MTA and HC volume were stronger than previously reported, with r_s_ ∈ [-0.26,-0.37] (Wahlund et al., 1999; Cavallin et al., 2012a). This shows that both radiologists are reliable, but that their rating styles differ—with one being more conservative—leading to low agreements. Since none of the radiologists trained together prior to rating the images, the low *κ_w_* is not surprising. These results demonstrate the issue of using subjective measures to quantify atrophy, where pathological status (normal/abnormal MTA) of a patient may differ depending on which radiologist performs the rating. On the other hand, 33 images failed the FreeSurfer segmentation upon visual QC. Having to discard almost 10% of the MRI scans due to software issues is not acceptable in a clinical setting. While other segmentation tools may be more reliable than FreeSurfer, inter-scanner variability, scanner software updates and image artifacts will always be obstacles that can influence performance (Guo et al., 2019; Mårtensson et al., 2019b). This does not seem to be an issue for visual ratings, where excellent intra-rater agreement has been demonstrated even across modalities (Wattjes et al., 2009). The benefits of using objective measures will outweigh the disadvantages—particularly as softwares become more robust—but it is important to understand that a software may fail in other ways than humans.

In accordance with previous studies we found increased HC (and ILV) atrophy rates with progressed CSF AD pathology and in MCI patients compared to cognitively normal (CN) subjects (Rusinek et al., 2003; Ridha et al., 2006; Henneman et al., 2009a). Pettigrew et al. (2017) specifically investigated the progression of MTL atrophy in preclinical AD, defined by abnormality in amyloid-β and tau. They also found an increased atrophy rate in individuals with A^+^ T^+^ biomarker profile. They did not find any differences between A^−^T^−^ and A^+^T^−^. We observed differences in our automatic measures, although these differences where smaller when studying SCD subjects only. Pettigrew and colleagues investigated only CN subjects that were 10-15 years younger (on average) than in our study. Further, we defined CSF abnormality based on established cut-offs, and not by percentiles of the sample distribution. We expect our study sample to be in a more advanced pathological stage, which may explain why our data showed a difference between A^−^T^−^ and A^+^T^−^. Henneman et al. (2009a) reported differences in both HC volume at baseline and HC atrophy rate between healthy controls and MCI patients, which is consistent with our observed differences between SCD and MCI subjects within each CSF group. However, SCD individuals in the A^+^T^+^ group displayed greater atrophy rates than A^−^T^−^ and A^+^T^−^ MCI patients. This is in line with another study from Henneman et al. (2009b) which suggested that greater CSF p-tau levels were associated with greater HC atrophy rate.

The same trends as for HC were captured by AVRA’s MTA ratings, but not as clear in the radiologist ratings. Most subjects, when using discrete ratings, had the same or +1 MTA score at six-year follow-up compared to baseline. This led to that the computed ΔMTA/year values for Rad. 1 and Rad. 2 merely reflect the ratio of subjects given a higher MTA score within six years. That is, the Δ MTA/year for a subject can “only” assume three values {0, 0.15, 0.2} depending on if, or at what timepoint, a higher MTA score is assigned. Thus, we argue that it is not possible to obtain reliable measure of atrophy rates from the integer radiologist ratings in our small study samples. Focusing on the ratings from AVRA only, we found that the average changes in MTA scores were small: between 0.04 and 0.15 per year. This corresponds to roughly 25 years for A^−^T^−^ subjects to progress a “full” MTA score (e.g. “1.0 → 2.0”). For the A^+^T^−^ group the time is 13.3 years, and 8.3 years for A^+^T^+^. By combining the ΔHC/year entries from Table 3 with the ΔHC value at which it becomes more likely for the radiologists to give a higher MTA score (Fig. 4), we can estimate how many years it takes for individuals in each CSF group to be more likely to get a higher MTA score at follow-up. Subjects with A^−^T^−^ at baseline are more likely to get a higher score at roughly 6.2 years, A^+^T^−^ at 4.3 years, and A^+^T^+^ at 2.5 years. The difference in the two methods is that in the latter measure we are estimating the time to reach the next discrete MTA step. That is, borderline cases (e.g. MTA=2.9) are more likely to get a higher score at the next follow-up than individuals with MTA=2.0 at baseline. Assuming that patients being rated MTA=2 by a radiologist have an underlying continuous MTA score, and that these are uniformly distributed on the interval [2,3), a patient in this group would on average have MTA=2.5. The first method (based on AVRA ratings) should thus give roughly twice the conversion time to the second (based on radiologist ratings), which is too short but fairly close. The remaining differences can have multiple explanations. 1) The estimates are crude and based on relatively few subjects with large within-group variability in MTA rates. 2) The calculations are based on atrophy rates being constant over 20 years. This seems unlikely, given that individuals’ CSF status and cognition may worsen, which should yield increased atrophy rates according to Table 3. 3) The MTA scale assesses three structures and not just HC atrophy. Further, it has been suggested that atrophy mainly occurs in posterior HC in preclinical AD (Lindberg et al., 2017), leading the HC volume change to occur mainly “outside” the MTA rating slice.

Longitudinal MTA scores have, to our knowledge, only previously been reported by Ferreira et al. (2017) in AD patients and CN subjects over a two-year follow-up. This study reported an an MTA change of 0.25/year in CN participants, and 0.4/year in AD patients (estimated from figure). The annual change in MTA scores in CN individuals was higher than those observed in SCD subjects in the current study. However, we believe that our data, comprising four scans per participant and continuous ratings, allows for an accurate estimation of the MTA rate.

A limitation of the current study is that many of the analyses assume a linear relationship between variables. From Fig. 3 the individual slopes for all MTL measures look linear with respect to age, or at least like a reasonable approximation for six years. However, if one was to model ILV as a function of MTA (see Fig. 2), the relationship is clearly not linear. This means that the change in ILV volume between MTA 0-1 is smaller than between MTA 3-4. This may confound the interpretations of Fig. 4, but our study sample was not large enough to consider non-linear relationships. Further, we emphasize that the study sample is not fully representative of A^−^T^−^, A^+^T^−^ and A^+^T^+^ groups given that the inclusion criteria excluded (subjective) cognitively normal and dementia patient. The former would likely affect mainly the A^−^ group results, and the latter the CSF pathological groups.

## 5 Conclusion

In this study we investigated the sensitivity and reliability of visual assessment of MTL atrophy according to Scheltens’ MTA scale in a longitudinal cohort of subjects with subjective cognitive decline and mild cognitive impairment. Our data showed that MTA ratings display the same cross-sectional and longitudinal trends as the volumes of hippocampus and the inferior lateral ventricle, suggesting that the MTA scale is a sensitive alternative to automatic image segmentations. The MTA ratings from two experienced radiologists, and an automated software, were strongly associated to the subcortical volumes as well as cognitive tests, showing that all raters were reliable. However, the inter-rater agreement was low due to systematic rating differences, which highlights the issue of using subjective assessments.

## 6 Acknowledgements

The authors wish to thank Konstantinos Poulakis for helpful discussions regarding the statistics of this paper. Further, we thank the Swedish Foundation for Strategic Research (SSF), The Swedish Research Council (VR), the Strategic Research Programme in Neuroscience at Karolinska Institutet (StratNeuro), Swedish Brain Power, Center for Innovative Medicine (CIMED), the regional agreement on medical training and clinical research (ALF) between Stockholm County Council and Karolinska Institutet, Swedish Brain Foundation, Swedish Alzheimer Foundation, Åke Wiberg Foundation, Stiftelsen Olle Engkvist Byggmästare, the joint research funds of KTH Royal Institute of Technology and Stockholm County Council (HMT), and Birgitta och Sten Westerberg for financial support. The Titan X Pascal used for this research was donated by the NVIDIA Corporation.

## A Supplementary data

As additional information we provide visual examples of the MTA rating slice for four subjects in Fig. S1. Figure S2 is the same plot as Fig. 3 but for the right hemisphere. Confusion matrices for all rating sets are shown in Table S1–S3.

**Table S1:**
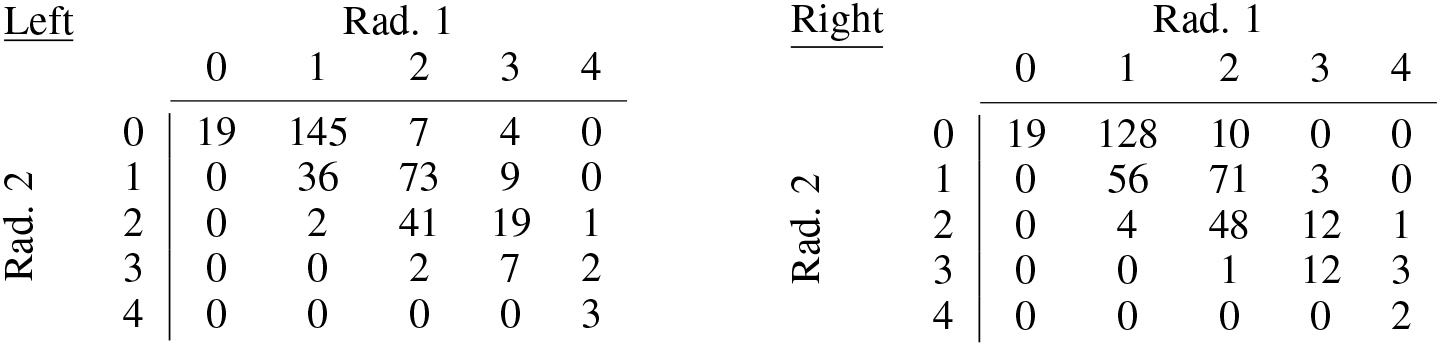
Confusion matrices for left and right MTA ratings between Rad. 1 and Rad. 2.

**Table S2:** Confusion matrices for left and right MTA ratings between Rad. 1 and AVRA.

| <u>Left</u> |  | Rad. 1 |  |  |  |  | <u>Right</u> |  | Rad. 1 |  |  |  |  |
| --- | --- | --- | --- | --- | --- | --- | --- | --- | --- | --- | --- | --- | --- |
|  |  | 0 | 1 | 2 | 3 | 4 |  |  | 0 | 1 | 2 | 3 | 4 |
| AVRA | 0 | 1 | 5 | 0 | 0 | 0 | AVRA | 0 | 1 | 6 | 1 | 0 | 0 |
|  | 1 | 18 | 158 | 26 | 5 | 0 |  | 1 | 18 | 157 | 28 | 0 | 0 |
|  | 2 | 0 | 19 | 83 | 19 | 1 |  | 2 | 0 | 24 | 85 | 10 | 0 |
|  | 3 | 0 | 1 | 14 | 13 | 5 |  | 3 | 0 | 1 | 16 | 17 | 5 |
|  | 4 | 0 | 0 | 0 | 2 | 0 |  | 4 | 0 | 0 | 0 | 0 | 1 |

**Table S3:** Confusion matrices for left and right MTA ratings between Rad. 2 and AVRA.

| <u>Left</u> |  | Rad. 2 |  |  |  |  | <u>Right</u> |  | Rad. 2 |  |  |  |  |
| --- | --- | --- | --- | --- | --- | --- | --- | --- | --- | --- | --- | --- | --- |
|  |  | 0 | 1 | 2 | 3 | 4 |  |  | 0 | 1 | 2 | 3 | 4 |
| AVRA | 0 | 6 | 0 | 0 | 0 | 0 | AVRA | 0 | 7 | 1 | 0 | 0 | 0 |
|  | 1 | 162 | 44 | 1 | 0 | 0 |  | 1 | 143 | 60 | 0 | 0 | 0 |
|  | 2 | 7 | 74 | 41 | 0 | 0 |  | 2 | 7 | 66 | 45 | 1 | 0 |
|  | 3 | 0 | 0 | 20 | 10 | 3 |  | 3 | 0 | 3 | 20 | 15 | 1 |
|  | 4 | 0 | 0 | 1 | 1 | 0 |  | 4 | 0 | 0 | 0 | 0 | 1 |

**Figure S1:**
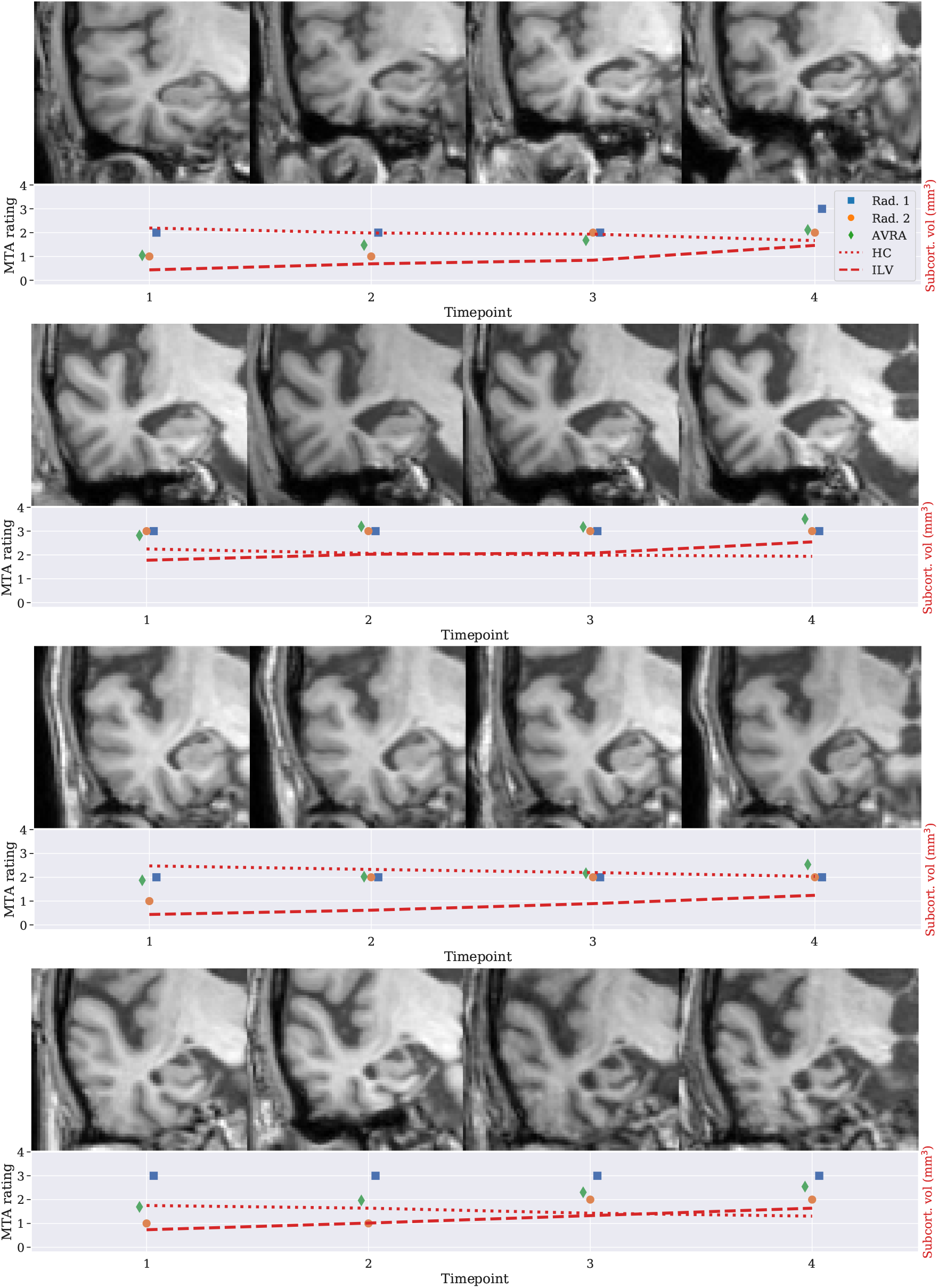
Rating slices at each timepoint for study four participants and corresponding MTA ratings and MTL volumes.)

**Figure S2:**
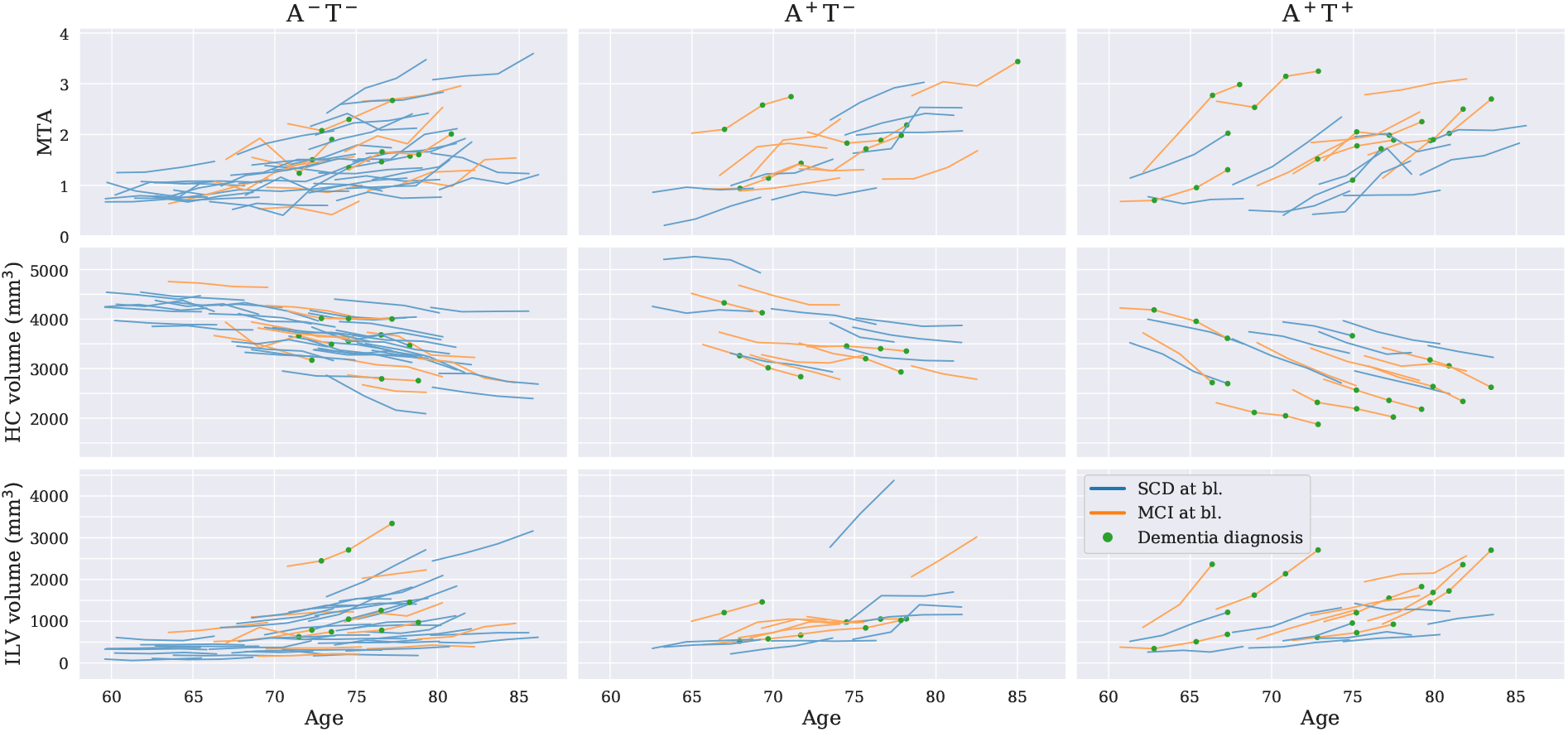
*Top:* AVRA’s **right** MTA ratings plotted against age at scan time for different combinations of A*β* and p-tau abnormality. *Middle*/*bottom:* corresponding plots for right hippocampal (HC)/inferior lateral ventricles (ILV) volumes. Orange and blue lines show individual trajectories for SCD and MCI patients, respectively. The green dots show if a patient was diagnosed with dementia at the given timepoint.

## Footnotes

1 *Automatic visual Ratings of Atrophy*, freely available at https://github.com/gsmartensson/avra_public

2 Freely available at http://surfer.nmr.mgh.harvard.edu/

